# Circular RNA profiling and functional screening in breast cancer identify circNSD1(6) as a suppressor of tumor autophagy

**DOI:** 10.1101/2025.03.24.645059

**Authors:** Wenhui Li, Wen Yang, Peter Hyunwuk Her, Tinglin Yang, Tiantian Li, Ziwei Huang, Xin Xu, Mona Teng, Moliang Chen, Fraser Soares, Stanley Liu, Benjamin Haibe-Kains, Housheng Hansen He, Jie Ming

## Abstract

Circular RNAs (circRNAs) have emerged as critical regulators of cell biology. However, their function in breast cancer remains elusive. Herein, through circRNA profiling of 38 breast tumors and 10 benign breast tissues, we identified 509 of differentially expressed circRNAs. Integration with transcriptome-wide functional screening of approximately 10,000 circRNAs pinpointed circNSD1(6) as a top ranked tumor suppressor in breast cancer. circNSD1(6) is downregulated in tumors and suppression of the circular form, but not the linear form, promotes breast cancer proliferation and tumor growth. Mechanistically, circNSD1(6) interacts with autophagy receptor SQSTM1/p62 and inhibits the oligomerization of p62, thereby suppressing p62 body formation and p62-dependent autophagy. Furthermore, circNSD1(6) ameliorates p62-mediated Keap1 sequestration, hence suppresses the Nrf2 pathway and oxidative stress tolerance. Our study provides a landscape view of the transcription and function of circRNA in breast cancer, and uncovers the crucial role of circNSD1(6) in autophagy and antioxidant stress.

## Introduction

Breast cancer (BC) is the most common malignancy worldwide and the leading cause of malignancy-related mortality in women^1^. Despite the improvement of diagnostic and therapeutic intervention, breast cancer remains life-threatening due to drug resistance, recurrence, and metastasis. A deeper understanding of molecular mechanisms underlying breast cancer development and progression holds promise for the identification of novel therapeutic targets^2^.

Circular RNA (circRNA) is a class of RNA produced from non-canonical back-splicing events of precursor mRNA^3^. Advances in high-throughput RNA sequencing and bioinformatics tools have boosted the discovery of hundreds of circRNAs, exhibiting both tissue-specific and cancer-specific expression patterns. Featured by the special covalently closed circular structure, accumulating evidence indicated that circRNAs exert unique properties and specific biological functions in tumor initiation and progression. The circular structure also facilitates circRNAs with stability, resistance to exonuclease degradation, and detectability in human body fluids and exosomes, highlighting its potential as biomarkers in non-invasive liquid biopsies. Despite numerous circRNAs being found, only a limited number of circRNA have been elucidated with biological functions involved in breast cancer initiation and progression. Therefore, further investigation and functional exploration of circRNAs in mammary tissue will provide a novel perspective into comprehending tumorigenesis and molecular targets^4,5^.

SQSTM1/p62 is a selective autophagy receptor, responsible for recruiting the aggregation of ubiquitin-tagged protein to autophagosomes for degradation. The p62 is capable of recognizing the ubiquitinated cargos via the UBA (Ub-associated) domain. Additionally, its capacity to self-polymerize via the PB1 domain aids in the formation of protein aggregates. Subsequently, p62 interacts with LC3B via the LC3-interacting region (LIR), facilitating the delivery of protein aggregates for autophagic degradation. Thus, p62 plays a central role in the assembly and degradation of protein aggregates. The p62 also serves as a stress-induced scaffold protein that helps cells to cope with oxidative stress through Kelch-like ECH-associated protein 1 (Keap1)-Nuclear factor erythroid 2-related factor 2 (Nrf2) pathway^6,7^. Nrf2 is a transcriptional factor that maintains redox balance and protects cells against oxidative stress-induced death by activating the expression of antioxidant enzymes. Sustained Nrf2 activation is commonly noted in many cancers, including breast cancer, facilitating cancer cells to overcome growth stress and promoting cancer progression^8,9^. Keap1 functions as a suppressor of Nrf2 by interacting with Nrf2 and mediating its ubiquitination and subsequent proteasomal degradation. Studies have demonstrated the involvement of p62 in the positive regulation of Nrf2 by interfering with Nrf2-Keap1 interaction and promoting Keap1 sequestration and degradation, with consequent translocation of Nrf2 to the nucleus and activation of cytoprotective and antioxidant genes^10,11^. Despite the critical role p62 plays in counteracting cellular stress by bridging autophagy and the Nrf2-Keap1 signaling pathway, there has been no previous report on circRNA regulation of p62.

In this study, we characterized a novel downregulated circRNA, circNSD1(6), by RNA-sequencing in breast cancer tissues and validated its suppressive role in tumor progression. Downregulation of circNSD1(6) significantly promoted the malignant potential of breast cancer cells in vitro and in vivo. Our study revealed that circNSD1(6) could interact with p62 and inhibit the oligomerization of p62, hence suppressing p62 puncta formation and p62-dependent autophagy. Moreover, circNSD1(6) relieved p62-mediated Keap1 sequestration, thereby decreasing Nrf2-mediated transcriptional activation of antioxidant genes. Our finding unveiled the suppressive role of circNSD1(6) in autophagy and Nrf2 activation, resulting in attenuated tumorigenesis, suggesting the diagnostic and therapeutic potential circNSD1(6) possessed in breast cancer.

## Results

### Circular RNA profiling and functional screen in breast cancer

To assess the transcriptome of breast cancer, we assembled a cohort of 38 breast cancer tissues including 10 luminal A (lum A), 11 luminal B (lum B), 6 human epidermal growth factor receptor 2 (HER2-positive), and 11 triple-negative breast cancer (TNBC) subtypes and 10 normal breast tissues. RNA of each sample was quantified via strand-specific, ribosomal RNA-depleted total RNA-seq **(Fig 1. A, right panel)**. This allows for the rare identification of both non-coding and coding transcripts. Breast cancer is recognized as a heterogeneous disease, comprising multiple molecular subtypes with distinct biological characteristics. To verify these transcriptomic subtypes, we applied unsupervised clustering to known breast cancer subtype-specific transcripts **(Fig 1. B)**. Luminal A tumors are typically characterized by the expression of estrogen receptor (ESR1) and progesterone receptor (PGR) with low proliferation rates, as indicated by low MKI67 levels. In contrast, Luminal B tumors, while also ESR1 and PGR positive, exhibit higher proliferation as indicated by the elevated MKI67 compared to Luminal A. HER2-enriched tumors are distinct in their overexpression of ERBB2. TNBC, defined by the absence of ER, PR, and ERBB2, is associated with the highest levels of MKI67, in line with its highly proliferative and aggressive nature (**Supplementary Fig. S1A**). Additionally, PAM50 expression profiles were employed to classify and verify the breast cancer subtypes into their respective molecular categories (**Supplementary Fig. S1B**).

**Figure 1.**
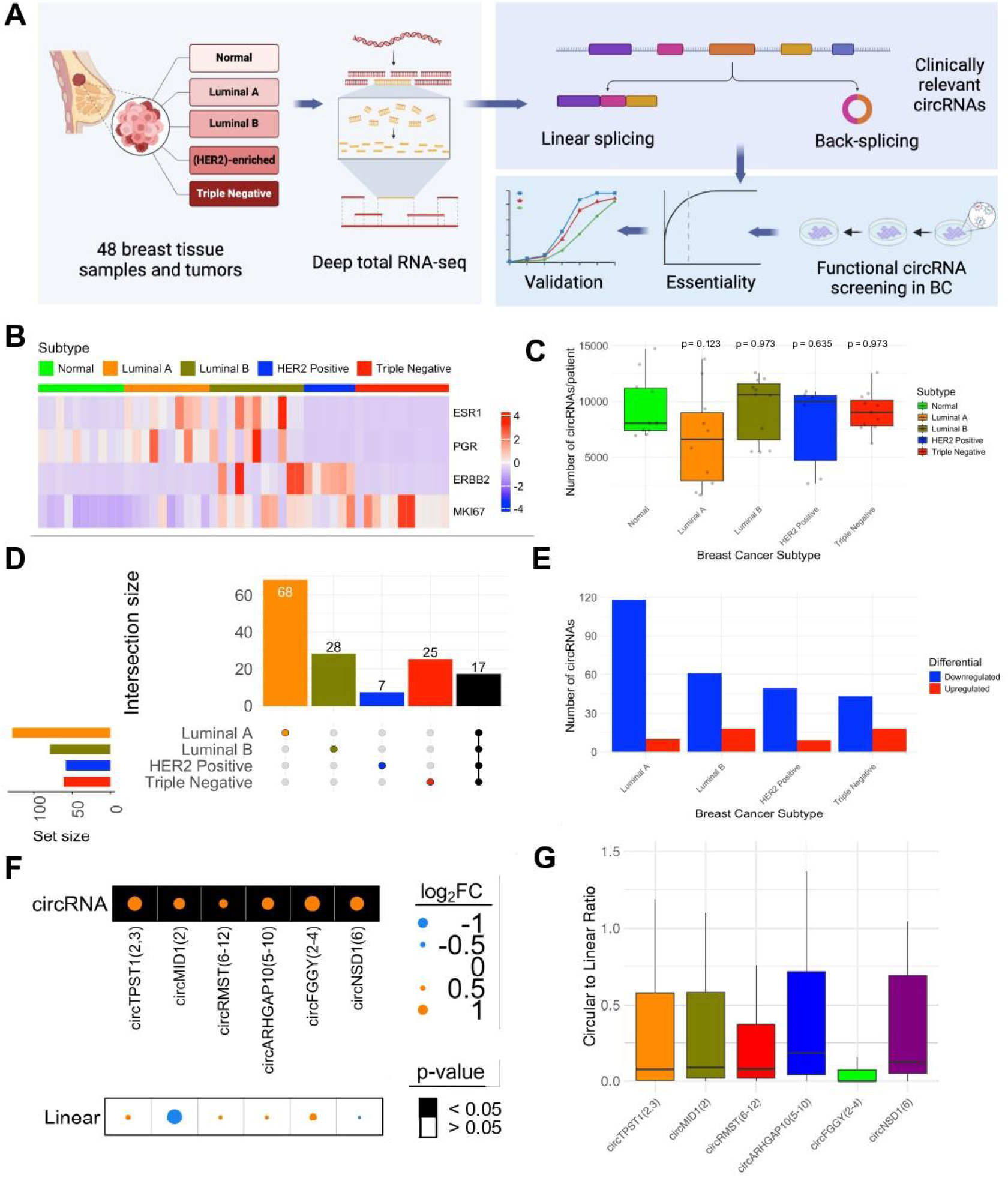
Circular RNA profiling and functional screen in breast cancer. (A) Schematic representation of workflow. The study involves 48 breast tissue samples, including normal tissue along with luminal A, luminal B, HER2-enriched, and triple negative subtypes. Deep total RNA-sequencing was performed to capture both linear splicing and back-splicing events, identifying clinically relevant circRNAs. Integration of these circRNAs with functional circRNA screening in breast cancer was used to pinpoint a target for further validation by various functional assays. (B) Heatmap showing the expression levels of key breast cancer genes (ESR1, PGR, ERBB2) and the proliferation marker MKI67 across breast cancer subtypes. Expression is represented by z-score. (C) Boxplot displaying the number of non-zero circRNAs per patient across breast cancer subtypes. The y-axis shows the number of circRNAs identified for each patient, with each point representing an individual patient. Mann-Whitney U tests between normal and the subtypes in order (Luminal A, Luminal B, HER2 Positive and Triple Negative) are 0.123, 0.973, 0.635 and 0.973. (D) UpSet plot of differential circRNAs. Set size bar graph on the left corner represents the total number of circRNAs present for each subtype. Columns represent each subtype, with the number of subtype-specific or shared circRNAs. Absolute log_2_ fold change greater than 1 and bonferroni adjusted p-value less than 0.01. (E) Bar plot showing the number of up and downregulated circular RNAs (circRNAs) across breast cancer subtypes. The y-axis represents the number of circRNAs, with blue bars indicating downregulated circRNAs and red bars indicating upregulated circRNAs. Log_2_ fold change greater than 1 (for upregulated) and less than −1 (for downregulated) and bonferroni adjusted p-value less than 0.01. (F) Illustration of proliferation of enriched circRNAs from MCF7 screen that overlapped with differential circRNAs in luminal A (Log_2_ fold change less than −1 and bonferroni adjusted p-value less than 0.01). Background shading indicates p value; size and color of dot shows log_2_(FC). (G) Circular to linear ratio of overlapping circRNAs from MCF7 screen and differential circRNAs in luminal A across samples

We then applied the CIRCExplorer pipeline to identify and quantitate the expression circRNAs in the breast tissue samples^12^. We identified a total of 47,759 circRNAs in the 48 samples, with a median of 8,727 circRNAs per sample **(Supplementary Table S1)**. The distribution of circRNA numbers across the breast cancer subtypes varied **(Fig 1. C)**. Notably, HER2-positive and Luminal B subtypes showed the highest median circRNA number per patient, followed by Normal and Triple Negative subtypes. The Luminal A subtype exhibited the lowest median circRNA number of 6,602, indicating a potential subtype-specific regulation of circRNA biogenesis or stability.

To characterize whether the expression patterns of circRNAs varied between tumor and normal samples, we performed differential abundance analysis for circRNAs between subtypes and normal breast tissues. We identified a total of 519 common differentially expressed circRNAs in normal and tumor samples, with Luminal A exhibiting the highest number of unique differential circRNAs **(Fig 1. D)**. Each subtype had a higher number of downregulated circRNAs, with Luminal A exhibiting the highest number of downregulated circRNAs compared to other subtypes and relatively fewer upregulated circRNAs (**Fig. 1E**).

To identify functional circRNAs in breast cancer, we performed shRNA functional screening against ∼10,000 circRNAs and their counterpart linear transcripts in MCF7 cells, a Luminal A breast cancer cell line. Analysis of the screening data identified 502 and 362 circRNAs with targeting shRNAs that were either enriched or depleted during the screening (**Supplementary Fig. S1C**). We next integrated the functional screening data with differentially expressed circRNAs in patient tumors, to further pinpoint key functional circRNAs **(Fig 1. A, left panels)**. Among the significantly downregulated circRNAs in Luminal A further filtered for adjusted p-value and fold change, six overlapped with enriched circRNAs in MCF7 functional screens (**Fig. 1F Supplementary Fig. S1D**). Interestingly, none of the circRNAs linear parental genes were enriched or depleted in the screen. Amongst these, circNSD1(6), was ranked high as a circRNA exhibiting higher circular to linear ratio (**Fig. 1G)**. In the functional screen in MCF7, circNSD1(6) showed significant enrichment, similar to the positive control whereas the linear parental gene NSD1 did not **(Supplementary Fig. S1E, S1F**).

### Downregulation of circNSD1(6) in breast cancer

circNSD1(6), originating from the circularization of exon 6 of the NSD1 gene located on chr5, has a length of 2,560 base pairs. The conformation of its back-splice junction site was verified through Sanger sequencing (**Fig. 2A**). In various subtypes of breast cancer tissues, circNSD1(6) exhibited significant downregulation when compared to normal breast tissues (**Fig. 2B**), while the linear NSD1 showed no significant changes in expression or the functional screen (**Fig. 1F**, **Supplementary Fig. S1G**). Consistently, RT-qPCR analysis revealed a notably higher expression of circNSD1(6) in the normal human breast epithelial cell line MCF10A compared to breast cancer cell lines MCF7 (luminal A), T47D (luminal A), BT-549 (triple negative) and MDA-MB-231 (triple negative) (**Fig. 2C**).

**Figure 2.**
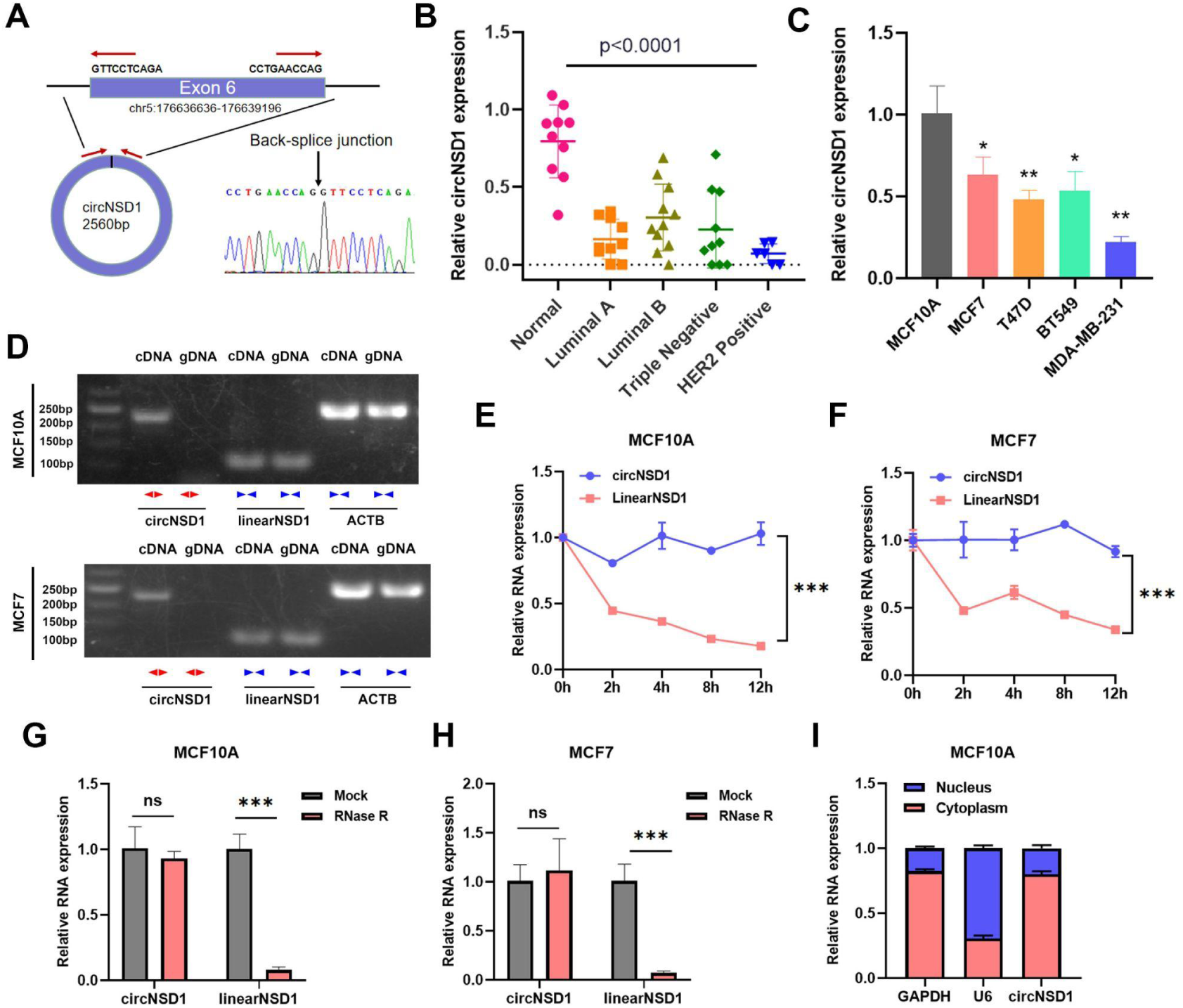
The identification and characteristics of circNSD1(6) in breast cancer. (A) The scheme depicting the genomic location and back-splicing of circNSD1(6). The back-splice junction site was verified by Sanger sequencing. (**B-C**) circNSD1(6) was downregulated in both breast tumor tissues and breast cancer cell lines. (**D**) The existence of circNSD1(6) from cDNA and gDNA in MCF-7 and MCF10A cells was detected by RT-qPCR with divergent and convergent primers. circNSD1(6) could only be amplified by divergent primers from cDNA but not gDNA. (**E-F**) MCF10A and MCF7 cells were treated with actinomycin D, and stronger stability was observed in circNSD1(6) compared with linear NSD1. (**G-H**) circNSD1(6) was proved to be resistant to RNase R digestion in MCF10A and MCF7 cells. (**I**) circNSD1(6) was mainly localized in the cytoplasm in the nuclear and cytoplasmic fractionation analysis.

To rule out the possibility of NSD1 genomic rearrangement, convergent primers for NSD1 mRNA and divergent primers for circNSD1(6) were designed. circNSD1(6) was successfully amplified using divergent primers using cDNA, but not in genomic DNA, confirming the presence of circularized NSD1 transcript (**Fig. 2D**). The covalently closed structure of circRNA may confer greater stability than linear RNA. To assess this, MCF10A and MCF7 cells were treated with the transcription inhibitor actinomycin D, revealing that circNSD1(6) transcripts exhibited greater stability compared to linear NSD1 transcripts (**Fig. 2E, F**). Additionally, circNSD1(6) resistance to RNase R digestion further supported its circular structure (**Fig. 2G, H**).

Since the function of circRNAs is often linked to their subcellular localization, we conducted nuclear and cytoplasmic fractionation analysis, which revealed the predominant localization of circNSD1(6) in the cytoplasm (**Fig. 2I**).

### circNSD1(6) suppresses breast cancer proliferation *in vitro* and *in vivo*

To investigate the functional role of circNSD1(6) in breast cancer, we designed two shRNAs to effectively suppress its expression in MCF10A and MCF7 cells, both of which exhibit high baseline levels of circNSD1(6). Importantly, these shRNAs did not significantly affect linear NSD1 expression (**Fig. 3A, B**). As revealed by CCK-8 assays, silencing circNSD1(6) significantly enhanced the growth of MCF10A and MCF7 cells (**Fig. 3C, D**). Specifically, knockdown of circNSD1(6) led to a higher proportion of proliferating cells, as confirmed by the EdU assay (**Fig. 3E-H**). Furthermore, the colony formation assay demonstrated increased colony formation in MCF10A and MCF7 cells following circNSD1(6) knockdown (**Fig. 3I-L**).

**Figure 3.**
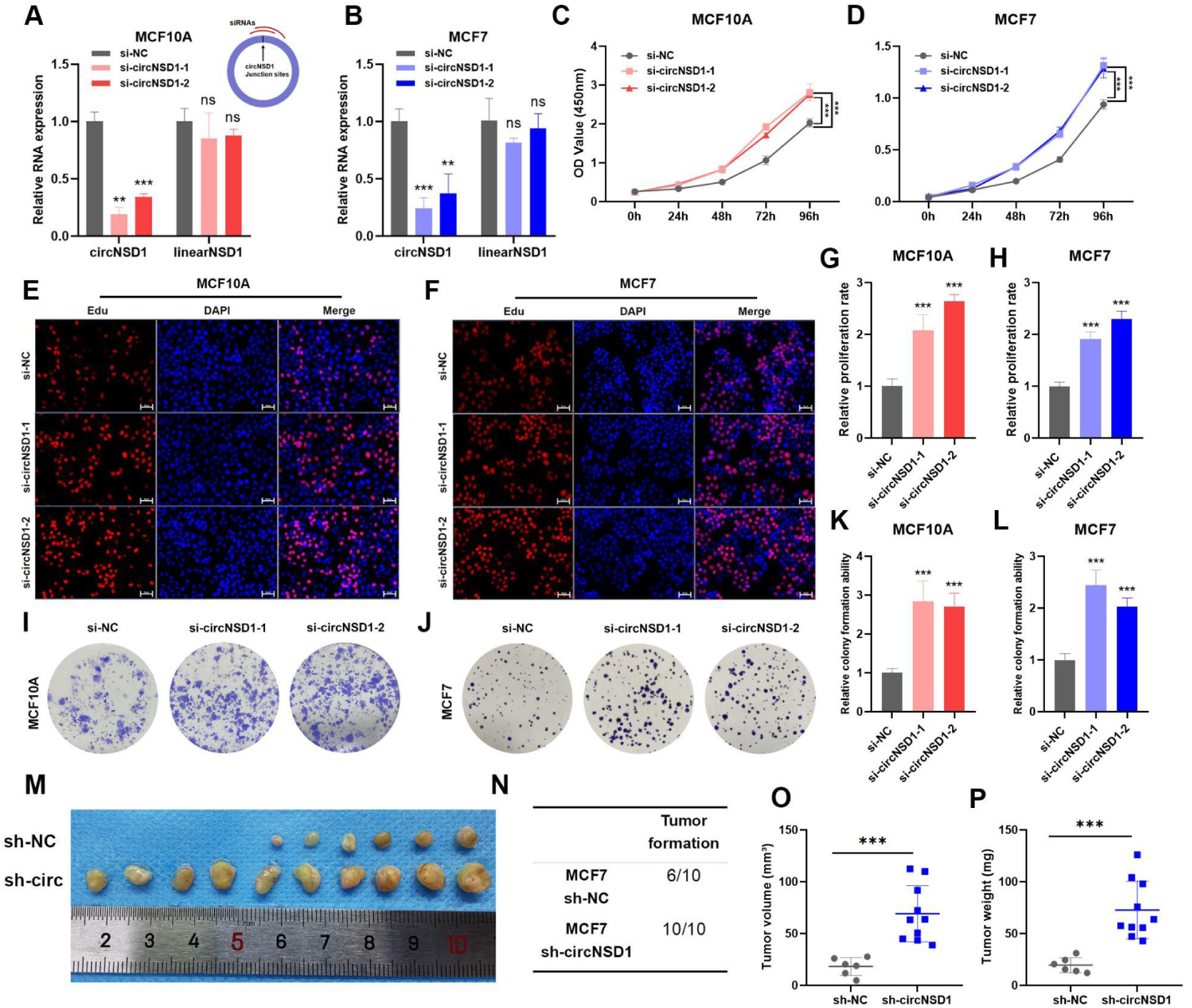
circNSD1(6) suppresses breast cancer proliferation in vitro and in vivo. (**A-B**) Two siRNAs effectively suppressed the expression of circNSD1(6) in MCF10A and MCF7 cells, while no significant impact on NSD1 expression was observed. (**C-D**) Enhanced viability of circNSD1(6)-knockdown MCF10A and MCF7 cells was detected by CCK-8 assays. (**E-F**) The knockdown of circNSD1(6) resulted in an increased proportion of proliferating MCF10A and MCF7 cells in EdU assays. (**G-H**) The statistical analysis of relative proliferation rates in EdU assays. (**I-J)** The knockdown of circNSD1(6) promoted the colony formation ability in MCF10A and MCF7 cells. (**K-L**) The statistical analysis of relative colony formation abilities of MCF10A and MCF7 cells. (M) circNSD1(6)-knockdown MCF7 cells exhibited greater tumorigenic capacity in in vivo assays conducted in nude mice. Increased numbers and larger volumes of tumors were formed in the sh-circNSD1(6) group, indicating that circNSD1(6) suppresses breast cancer proliferation in vivo. (N) More subcutaneous transplanted tumors were formed in the sh-circNSD1(6) group. (**O-P**) Increased tumor volumes and weights were observed in the sh-circNSD1(6) group.

To further assess the role of circNSD1(6) in breast cancer *in vivo*, control MCF7 cells or circNSD1(6)-knockdown MCF7 cells were subcutaneously injected into nude mice. Ontably, only six out of ten mice developed tumors after receiving injection of control cells, whereas all ten mice developed tumors after injection of circNSD1(6)-knockdown cells (**Fig. 3M, N**). Furthermore, the xenografts in the circNSD1(6) knockdown group exhibited significantly increased tumor volumes and weights compared to those in the control group (**Fig. 3O, P**). Taken together, these findings demonstrated that suppression of circNSD1(6) promotes breast cancer tumorigenesis both i*n vitro* and *in vivo*, suggesting the inhibitory role of circNSD1(6) in breast cancer.

### circNSD1(6) inhibition promotes Nrf2 activation and tolerance of oxidative stress

To further investigate the molecular mechanisms underlying the suppressive role of circNSD1(6) in breast cancer, we performed RNA-Seq on MCF10A cells in which circNSD1(6) was knocked down with two different siRNAs against circNSD1(6) or a control siRNA. Gene set enrichment analysis (GSEA) revealed enrichment of pathways related to detoxification and metabolism-related pathways in circNSD1(6)-knockdown cells, including detoxification of reactive oxygen species, cellular responses to stress, fatty acid metabolism, and metabolism of steroids (**Fig. 4A**). Moreover, the top upregulated genes, such as AKR1C2, AKR1C3, AKR1B10, GPX2, NQO1, SLC7A11 and SOD2, are well-known cytoprotective genes regulated by Nuclear factor-erythroid 2-related factor 2 (Nrf2) (**Fig. 4B**). Nrf2, encoded by NFE2L2, is a pivotal transcriptional regulator involved in the cellular response to oxidative, nutrient, and metabolic stress. RT-qPCR confirmed the upregulation of Nrf2 target genes AKR1C2, AKR1C, AKR1B10, GPX2, and SLC7A11 in circNSD1(6)-knockdown cells (**Fig. 4C, D**).

**Figure 4.**
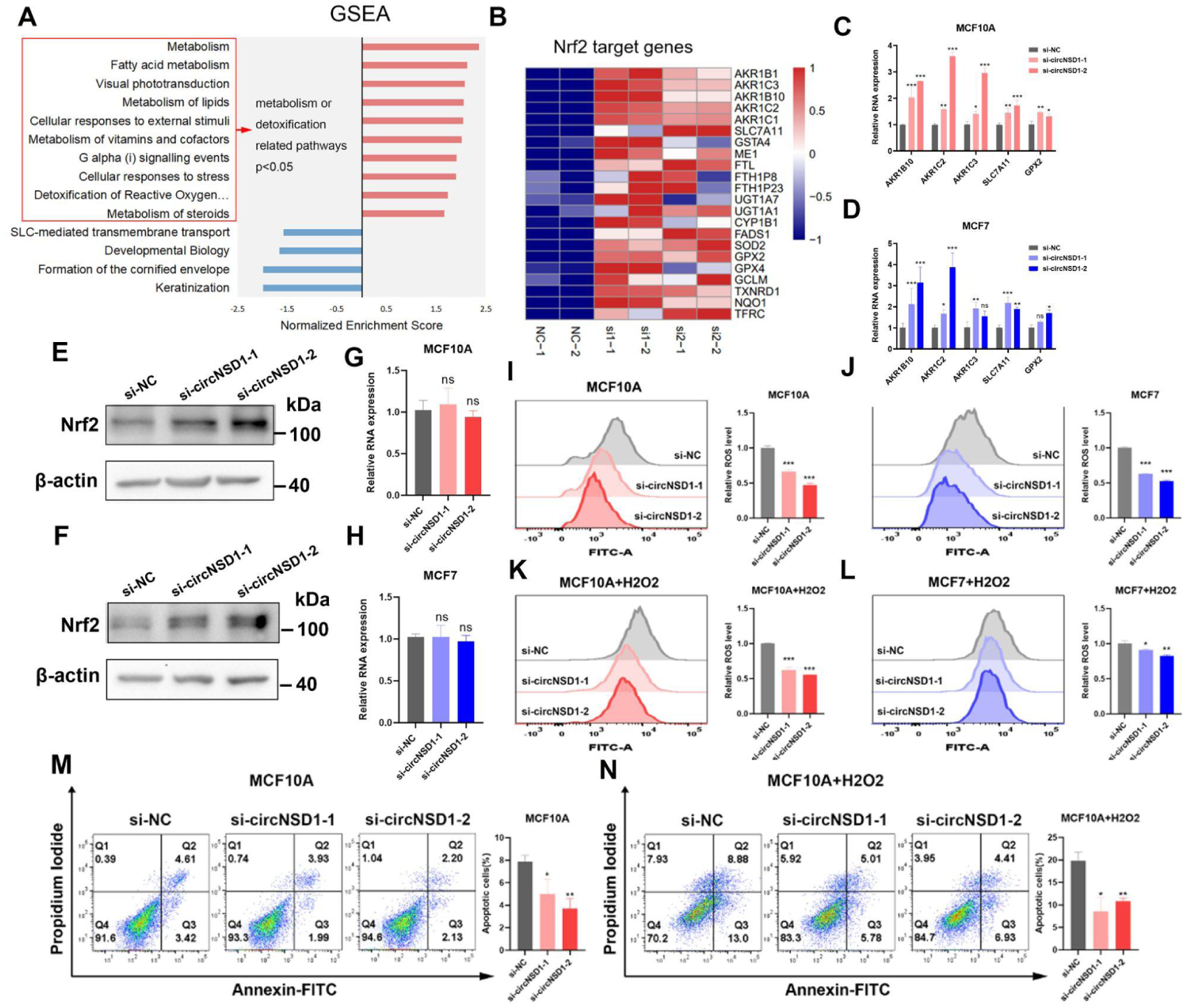
circNSD1(6) inhibition promotes Nrf2 activation and the tolerance of oxidative stress. (A) GSEA analysis indicated a positive enrichment of metabolism or detoxification-related pathways in circNSD1(6)-knockdown MCF10A cells. (B) Heatmap exhibiting the upregulated expressions of Nrf2-regulated cytoprotective genes in circNSD1(6)-knockdown MCF10A cells. **(C-D)** RT-qPCR validated the upregulation of Nrf2-related genes including AKR1B10, AKR1C2, SLKC7A11, and GPX2 at RNA level in circNSD1(6)-knockdown MCF10A cells and MCF7 cells. (**E-F**) As evidenced by western blotting, the knockdown of circNSD1(6) led to a significant increase in the abundance of Nrf2 protein, indicating the activation of Nrf2. (**G-H**) No significant change of Nrf2 was observed at the mRNA level after circNSD1(6) knockdown. (**I-J**) Intracellular ROS levels were reduced in circNSD1(6)-knockdown MCF10A and MCF7 cells detected by ROS-sensitive dye. (**K-L**) Intracellular ROS levels in response to H_2_O_2_ stimulation were reduced in circNSD1(6)-knockdown MCF10A and MCF7 cells. (**M)** Less apoptotic MCF10A cells were detected in the circNSD1(6)-knockdown group whether in response to H_2_O_2_ stimulation or not. (**N-O**) Larger numbers of live MCF10A and MCF7 cells were observed at the same concentration of H_2_O_2_ stimulation in the circNSD1(6)-knockdown group, demonstrating the increased tolerance of oxidative stress in circNSD1(6)-knockdown cells.

Based on these findings, we investigated whether Nrf2 mediates the observed changes in gene expression following circNSD1(6) inhibition. Consistent with this hypothesis, circNSD1(6) knockdown significantly increased the abundance of Nrf2 protein, as evidenced by western blotting and IF assays (Fig. 4E, F). However, there was no significant change in the transcription level of Nrf2 following circNSD1(6) knockdown (**Fig. 4G, H**).

Given the critical role Nrf2 plays in global cellular redox homeostasis, we further explored the impact of circNSD1(6) inhibition on oxidative stress defense. Using the ROS-sensitive dye 2ʹ,7ʹ-dichlorofluorescein diacetate (DCFH-DA), we measured intracellular reactive oxygen species (ROS) levels. Following exposure to hydrogen peroxide (H_2_O_2_)-induced stress, circNSD1(6) knockdown resulted in a significant reduction of intracellular ROS levels compared to the control cells (**Fig. 4I-L**). Notably, circNSD1(6) knockdown increased the live cell count and decreased cell death in response to H_2_O_2_ stimulation (**Fig. 4M-O**). These results indicate that circNSD1(6) knockdown activates the Nrf2-mediated cytoprotective program, enhancing the cell’s tolerance to oxidative stress.

### circNSD1(6) interacts with p62 protein

Upon revealing the important role of circNSD1(6) on Nrf2 signaling, we set out to investigate the underlying molecular mechanism. Since circRNAs often exert their functions by binding to specific proteins, we performed RNA pull-down assay in MCF10A cells using a biotin-labeled probe targeting the back-splicing site of circNSD1(6) or a non-targeting control probe. The RNA pull-down precipitates were subjected to SDS-PAGE gel analysis and silver staining, where the protein bands specific to the circNSD1(6) pulldown samples were excised and further analyzed using mass spectrometry (MS) ( **Fig. 5A**). We identified a total of 237 unique peptides that mapped to 115 proteins (**Supplementary Table S2**). Among these proteins, SQSTM1/p62 was the most abundant, with 14 unique peptides detected by MS (**Fig. 5B, C**).

**Figure 5.**
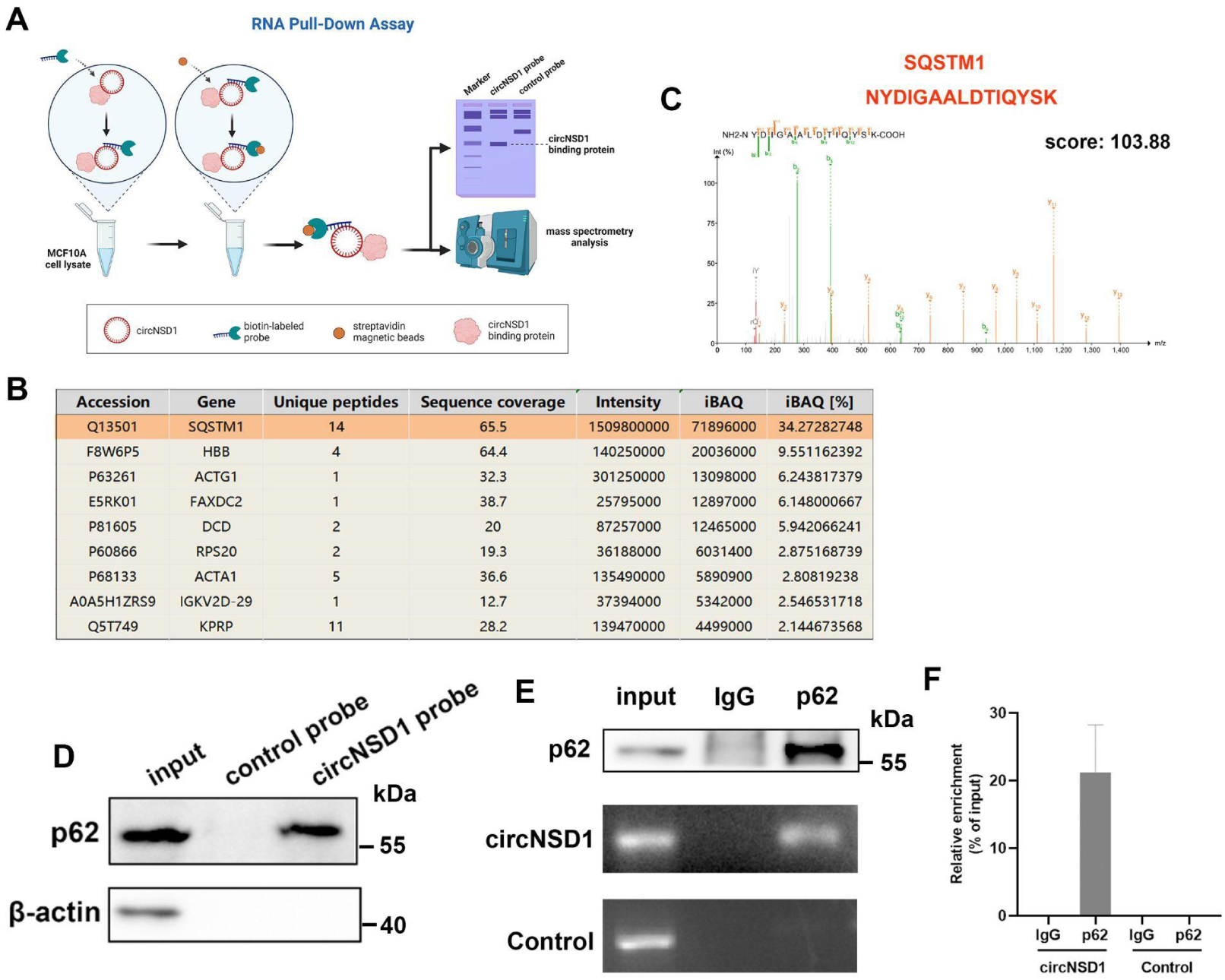
circNSD1(6) interacts with the autophagy receptor p62. (A) Potential circNSD1(6)-associated proteins identified via SDS-PAGE followed by silver staining. The circNSD1(6)-specific band at about 60 kDa (red box) was excised and analyzed by MS. (B) The identified proteins were ranked according to iBAQ(%), and the top 10 proteins were shown in the figure, among which SQSTM1 protein was absolutely dominant. (C) The mass spectrum of one of the 14 unique peptides of SQSTM1. (D) Interaction of p62 with circNSD1(6). β-actin was used as a negative control. (**E-F)** RIP assay was performed using the indicated antibodies and circACVR2A was used as a negative control.

Previous studies have reported that p62 interacts with Keap1, an inhibitor of Nrf2, via the SQSTM1 motif known as the Keap1-interacting region [KIR], thereby activating the Nrf2 antioxidant pathway^11^. Based on this, we hypothesized that circNSD1(6) might regulate the Nrf2 pathway through interacting with p62. To confirm the MS finding, we performed protein pull-down with the RNA probes followed by western blotting against p62, as well as RNA immunoprecipitation with p62 antibody followed by qPCR analysis of circNSD1(6), both of which validated the interaction between circNSD1(6) and p62 (**Fig. 5D, E, F**).

### circNSD1(6) knockdown enhances p62 expression and p62-dependent autophagy

Given that circNSD1(6) interacts with p62 to form an RNA-protein complex, we further explored whether circNSD1(6) regulates p62 abundance. We observed a significant increase in both protein and mRNA levels of p62 in the circNSD1(6)-knockdown cells (**Fig. 6A-C**). p62 is a well-established autophagy receptor protein. Given its pivotal role in autophagy regulation, we also measured the protein abundance of the autophagy marker protein LC3 and found that LC3 levels were significantly up-regulated following knockdown of circNSD1(6) (**Fig. 6A**).

**Figure 6.**
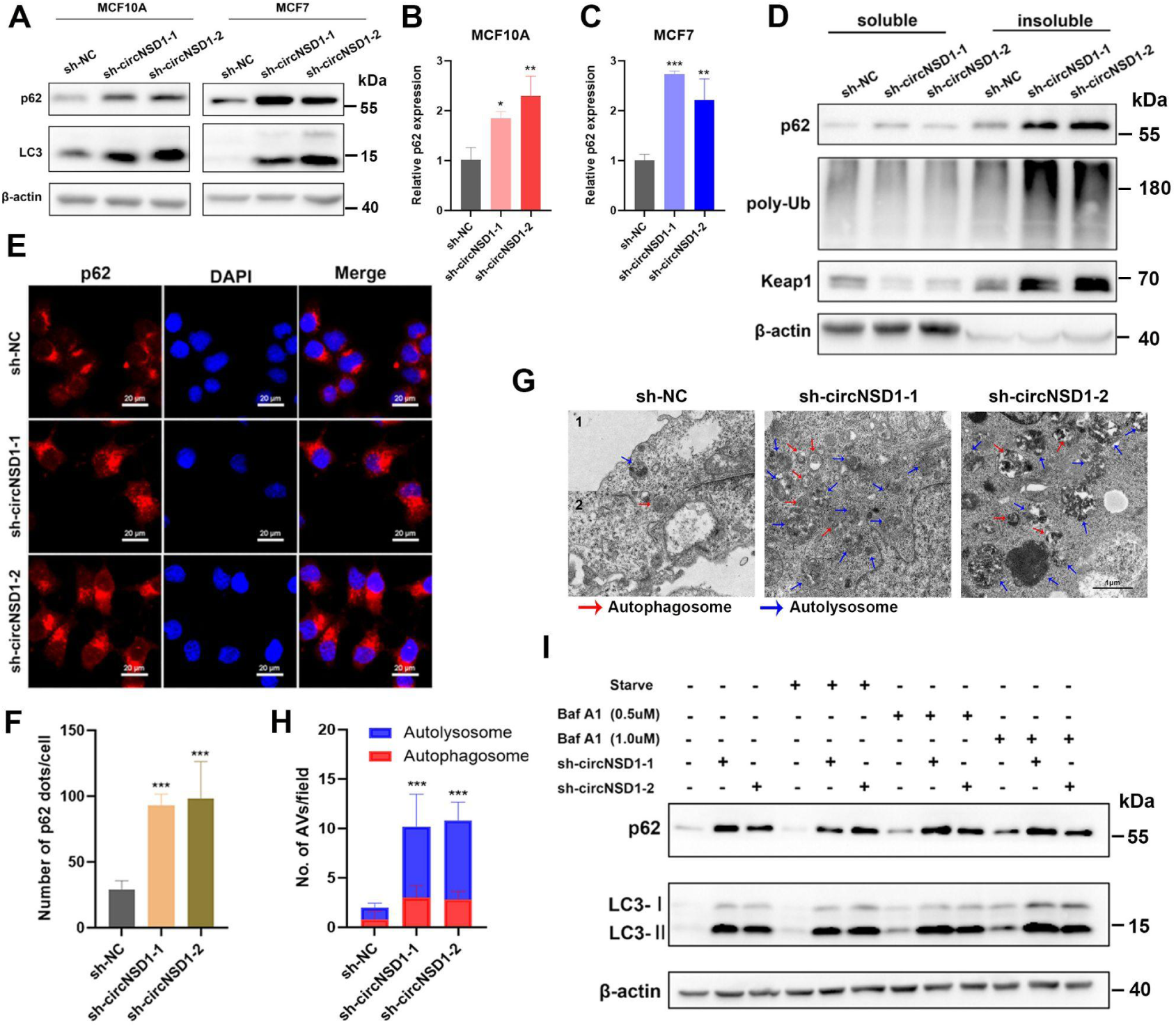
circNSD1(6) knockdown enhances p62 expression and p62-dependent autophagy. (A) p62 and the autophagy marker protein LC3 were both upregulated in circNSD1(6)-knockdown MCF7 and MCF10A cells. (**B-C**) mRNA levels of p62 were upregulated in circNSD1(6)-knockdown MCF7 and MCF10A cells. (**D)** Elevated expression levels of p62, poly-Ub, and Keap1 were detected in the insoluble fraction of circNSD1(6)-knockdown cells. (**E-F**) circNSD1(6)-knockdown cells formed increased numbers of cytoplasmic p62 bodies. (**G-H**) In the TEM observation, the number of autophagic vacuoles per cell remarkably increased in circNSD1(6)-knockdown cells observed with TEM. (**I**) Expression levels of p62 and LC3 in circNSD1(6)-knockdown and control cells under disposed condition, starvation, or Baf A1 treatment. The upregulation of p62 in the circNSD1(6)-knockdown group could not be reversed by Baf A1 treatment, indicating the elevated p62 expression was independent of autophagy.

It is well-known that various types of cellular stress induce the aggregation of ubiquitinated proteins and p62, leading to the formation of p62 bodies^13,14^. The formation of p62 bodies is critical for the activation of downstream signaling events associated with p62, including autophagy and its role in protein sequestration. Therefore, we investigated the potential role of circNSD1(6) in the homeostatic control of cytoplasmic p62 body levels.

Since p62 bodies are generally insoluble in non-ionic detergents like Triton X-100, we analyzed both the soluble and insoluble fractions of the cell lysate via western blotting. We found a substantial increase in insoluble p62 bodies, consisting of poly-ubiquitinated proteins and p62, in circNSD1(6)-knockdown cells compared to control cells (**Fig. 6D**). Furthermore, immunofluorescence analysis revealed that circNSD1(6)-knockdown cells formed substantially more cytoplasmic p62 bodies than control cells (**Fig. 6E, F**). Taken together, these data suggest that circNSD1(6) suppresses the abundance of p62 bodies.

To investigate whether the accumulation of p62 was due to blocked autophagic flux, we examined the number of autophagic vacuoles per cell using transmission electron microscopy (TEM). The number of autophagic vacuoles was markedly increased in circNSD1(6)-knockdown cells compared to control ones (**Fig. 6G, H**). Furthermore, Baf A1 treatment led to the accumulation of both p62 and lipidated LC3 (LC3II). However, the treatment did not fully eliminate the difference in p62 expression between the circNSD1(6)-knockdown and control groups (**Fig. 6I**). These findings indicate that circNSD1(6) functions as a negative regulator of autophagy, and the accumulation of p62 in circNSD1(6)-knockdown cells is not solely caused by blocked autophagic flux.

### circNSD1(6) knockdown activates Nrf2 signaling by sequestrating Keap1 in p62 bodies

p62 is known to activate the Nrf2 signaling pathway by sequestering Keap1 into p62 bodies. Upon its dissociation from Keap1, Nrf2 stabilizes and translocates to the nucleus, where it activates the transcription of antioxidant genes. Since circNSD1(6) knockdown leads to the accumulation of p62 bodies, we further investigated whether the constitutive activation of Nrf2 signaling in the absence of circNSD1(6) is dependent on p62. Co-immunoprecipitation (Co-IP) assay demonstrated that circNSD1(6) knockdown resulted in increased recruitment of Keap1 to p62 bodies, while reducing the interaction between Keap1 and Nrf2 (**Fig. 7A, B**). Additionally, in circNSD1(6) knockdown cells, the levels of detergent-soluble Keap1 significantly decreased, while detergent-insoluble Keap1 levels significantly increased (**Fig. 7A, B**), further confirming that circNSD1(6) knockdown promotes the recruitment of Keap1 to p62 bodies.

**Figure 7.**
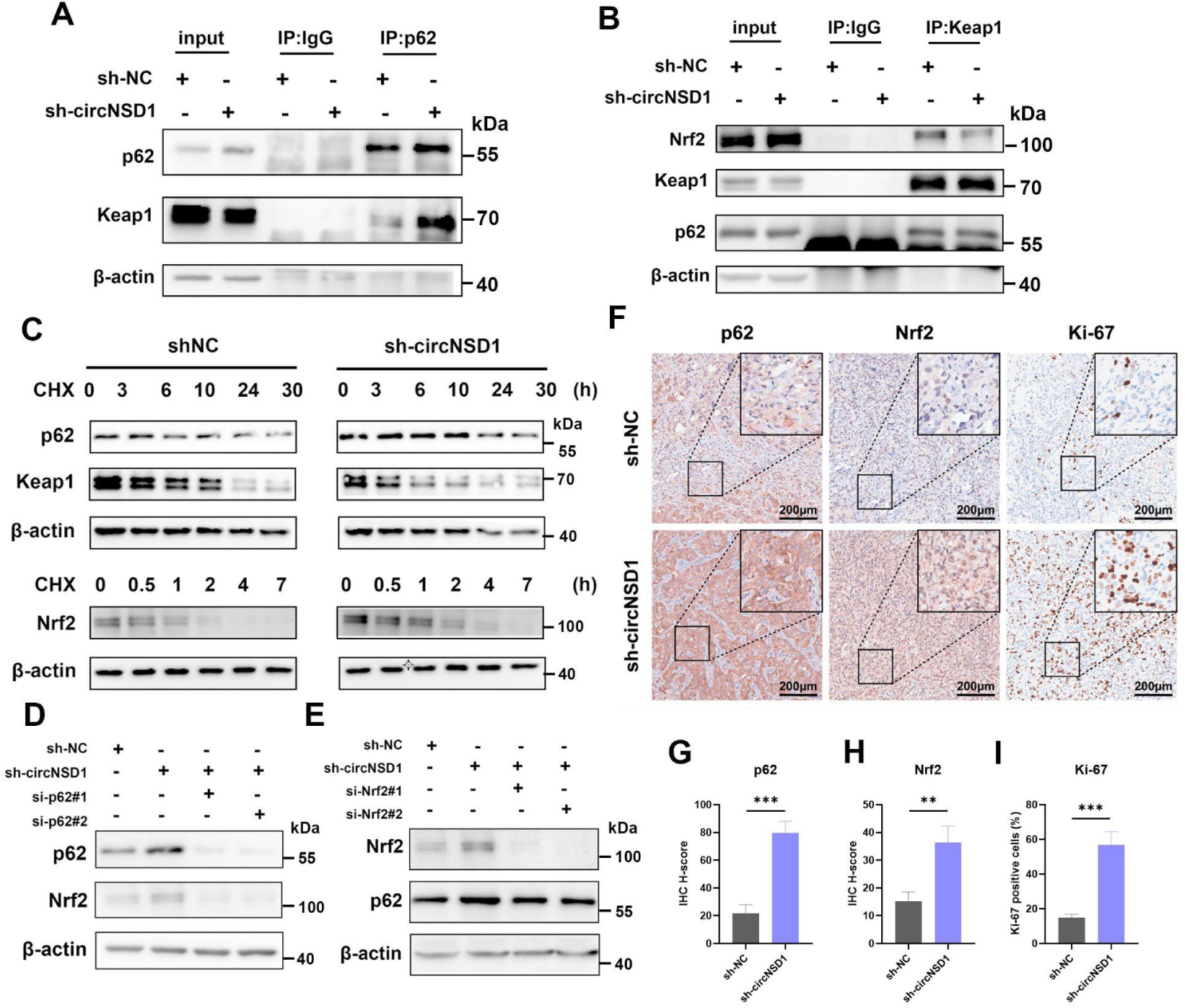
circNSD1(6) knockdown activates Nrf2 signaling by sequestrating Keap1 in p62 bodies. (**A-B**) circNSD1(6) knockdown led to an increase in the recruitment of Keap1 to p62 bodies revealed by the Co-IP assay, accompanied by a decreased interaction between Keap1 and Nrf2. (C) Cells were treated with the protein synthesis inhibitor CHX. Increased Nrf2 expression and decreased Keap1 expression were observed in circNSD1(6)-knockdown cells, demonstrating the knockdown of circNSD1(6) could stabilize Nrf2 expression and destabilize Keap1 expression. (D) The activation of Nrf2 caused by the knockdown of circNSD1(6) can be reversed by p62 inhibition. (E) The increase in p62 caused by the knockdown of circNSD1(6) can be partially reversed by Nrf2 inhibition. (F) Elevated expressions of p62, Nrf2, and Ki-67 were detected in the sh-circNSD1(6) group in IHC assays of xenograft tumor tissues in nude mice. **(G-I)** Statistical analysis of expression levels detected in the IHC assay.

To assess the role of circNSD1(6) in regulating Nrf2 protein stability, we treated cells with the protein synthesis inhibitor cycloheximide (CHX). circNSD1(6) knockdown stabilized Nrf2 protein abundance and destabilized Keap1 protein abundance (**Fig. 7C**). We further investigated whether circNSD1(6) activates Nrf2 in a p62-dependent manner. Knockdown p62 effectively blocked Nrf2 activation induced by circNSD1(6) downregulation (**Fig. 7D**).

SQSTM1, the gene encoding p62, is a transcriptional target of Nrf2^15^. In the RNA-seq analysis mentioned above, a significant increase in the mRNA level of SQSTM1 was observed in circNSD1(6)-knockdown cells compared to control cells, a finding confirmed by RT-qPCR. Furthermore, we revealed that Nrf2 silencing resulted in a decrease in p62 expression (**Fig. 7E**). These results suggest that p62 accumulation in circNSD1(6)-knockdown cells leads to Nrf2 activation, which in turn further upregulates p62 expression, forming a positive feedback loop.

To validate the circNSD1(6)/p62/Nrf2 axis in breast cancer, we examine the expression of p62 and Nrf2 in xenograft tumors in nude mice using IHC assay. Both p62 and Nrf2 were consistently upregulated in xenograft tissues with low circNSD1(6) expression (**Fig. 7F-H**). Additionally, the circNSD1(6)-knockdown cells had a higher proportion of Ki67 positive cells compared with the control cells (**Fig. 7F, I**).

Taken together, our results demonstrate that circNSD1(6) directly regulates the p62/Keap1/Nrf2 signaling by interacting with p62 and disrupting its ability to sequester Keap1, thereby negatively regulating the activity of Nrf2 (**Fig. 8**).

**Figure 8.** Graphic representation of mechanisms underlying the function of circNSD1(6) in regulating tumor autophagy.

## Discussion

Numerous circRNAs have been identified as dysregulated in various cancers and shown to play critical roles in tumorigenesis, metastasis, recurrence, and drug resistance. Due to their stability and low immunogenicity, circRNAs hold significant therapeutic potential in clinical practice.

To identify circRNAs involved in the pathogenesis of breast cancer, we performed rRNA-depleted RNA-seq analysis on 10 normal mammary tissues and 38 breast cancer tissues, representing the LumA, LumB, HER2-enriched, and TNBC subtypes from an asian cohort. Among these, we focused on circNSD1(6), a circular RNA that had not been previously studied. Our analysis revealed a significant downregulation of circNSD1(6) expression across all breast cancer subtypes. Furthermore, library screening showed that silencing circNSD1(6) promoted the proliferation of MCF7 cells, suggesting its key role as a tumor suppressor in breast cancer. In both in vitro and in vivo breast cancer models, we verified that circNSD1(6) inhibits cell proliferation and tumor formation. To elucidate the mechanisms behind the tumor-suppressive role of circNSD1(6), we conducted RNA-seq analysis on circNSD1(6)-knockdown MCF10A cells, which revealed significant changes in the expression of Nrf2 target genes. Subsequent experiments showed that circNSD1(6) binds to the autophagy receptor protein p62. This interaction prevents the formation of p62 bodies, thereby limiting p62-dependent autophagy and Keap1-Nrf2 signaling, ultimately inhibiting breast cancer progression.

Cancer cells develop effective ROS antioxidant systems to mitigate excessive ROS accumulation as an adaptive response to oxidative stress. Since many conventional cancer therapies, such as chemotherapy and radiotherapy, rely on increasing ROS levels to kill cancer cells, the presence of cytoprotective antioxidants presents a major challenge. Targeting molecules involved in ROS regulation could offer therapeutic opportunities to overcome treatment resistance. The transcription factor Nrf2 has been identified as a key regulator of cellular antioxidant responses, and increasing evidence suggests that the Nrf2 pathway contributes to cancer progression, metastasis, and drug resistance. Somatic mutations in Nrf2 or Keap1 that disrupt their interaction have been identified in various cancers, highlighting the importance of maintaining proper Keap1-Nrf2 interactions to limit Nrf2 activity and suppress cancer development. Our study demonstrates that silencing circNSD1(6) in mammary epithelial and breast cancer cells disrupts Keap1-Nrf2 binding, resulting in sustained Nrf2 activation and increased expression of metabolic and cytoprotective genes. When exposed to H2O2, circNSD1(6)-knockdown cells showed reduced ROS production and higher survival rates. Additionally, H2O2 treatment downregulated circNSD1(6) expression in breast cancer cells in a concentration-dependent manner, indicating a regulatory link between circNSD1(6) and redox homeostasis.

Studies have shown that specific circRNAs can act as protein sponges or decoys, influencing cellular functions or promoting protein-protein interactions as scaffolds. For instance, circPABPN1 serves as a decoy for HuR, impairing PABPN1 translation, while circVAMP3 binds to CAPRIN1, regulating its phase separation and promoting the formation of intracellular stress granules that inhibit liver cancer progression. In our study, we found that circNSD1(6) directly interacts with p62, significantly reducing p62 body formation. p62 is a multifunctional protein central to several signaling pathways, including the Keap1-Nrf2 pathway, suggesting that circNSD1(6) may regulate p62-mediated signaling. As expected, downregulation of circNSD1(6) promoted Keap1 sequestration and degradation, leading to sustained Nrf2 activation and enhanced p62-dependent autophagy.

Several mechanisms have been reported to regulate p62 body formation. For example, the interaction of MOAP-1 with the PB1-ZZ domain of p62 inhibits p62 self-oligomerization and liquid-liquid phase separation (LLPS), disrupting p62 body formation. In addition to interprotein interactions and post-translational modifications, recent studies have shown that p62 is an RNA-binding protein. For example, it interacts with vtRNA-1, a small noncoding RNA, which blocks p62 oligomerization and autophagosome formation. Our findings present a novel model in which a circRNA directly controls the function of p62 by acting as a protein sponge. p62 facilitates Keap1 sequestration and degradation, interfering with the Nrf2-Keap1 interaction and resulting in sustained Nrf2 activation. Since p62 is a target gene of Nrf2, this creates a positive feedback loop, where activated Nrf2 further enhances p62 expression. In this study, circNSD1(6) knockdown led to a substantial increase in both p62 mRNA and protein levels, further amplifying the p62-Nrf2 feedback loop.

In conclusion, our study demonstrates that the circular RNA circNSD1(6) plays a crucial role in regulating cellular redox homeostasis and autophagy by directly interacting with the autophagy receptor protein p62. Given the importance of autophagy and oxidative stress in cancer treatment, circNSD1(6) shows promise as a potential prognostic marker and therapeutic target in breast cancer. Our findings provide a foundation for developing circRNA-based therapeutic strategies for breast cancer management.

## Materials and Methods

### Tissue collection

A total of 38 breast cancer tissues including 10 luminal A (lum A), 11 luminal B (lum B), 6 human epidermal growth factor receptor 2 (HER2)-enriched, and 11 triple-negative breast cancer (TNBC) subtypes and 10 normal breast tissues were obtained from patients at the Department of Breast and Thyroid Surgery of Wuhan Union Hospital (Wuhan, China) between 2017 and 2018. Specimens were immediately frozen and stored in liquid nitrogen. Informed consent was obtained before collecting the tissues. This study was approved by the Wuhan Union Hospital Human Ethics Committee.

### Cell culture and in vitro treatments

The human breast cancer cell lines MCF7, T47D, and the immortalized human breast epithelial cell line MCF10A, were obtained from the American Type Culture Collection (Manassas, VA, USA). MCF7 cells were grown in Dulbecco’s modified Eagle’s medium (DMEM; Gibco, Carlsbad, CA, USA). T47D cells were cultured in RPMI 1640 medium (Gibco). MCF10A cells were cultured in DMEM/F12 supplemented with insulin, cholera toxin, epidermal growth factor (EGF), and hydrocortisone. Cells were cultured in the complete medium containing 10% fetal bovine serum (FBS; Gibco) in a 37°C, 5% CO2 humidified incubator.

For autophagy induction, cells were starved with Earle’s balanced salt solution (EBSS). Bafilomycin A1 (Baf A1) was used to inhibit autophagic flux. The proteasome inhibitor MG132 and cycloheximide (CHX) were administered to inhibit proteasomal degradation and protein synthesis, respectively. Hydrogen peroxide (H2O2) was added to observe the oxidative tolerance of cells.

### RNA extraction and real-time quantitative PCR (RT-qPCR)

Total RNA was extracted by Trizol (Vazyme, Nanjing, China), and HiScript III qRT SuperMix (Vazyme) was used for reverse transcription. Real-time quantitative PCR was performed using a BioRad CFX96 Real-Time PCR Detection System in triplicate. Fold change in gene expression was calculated by the ΔΔCt method, and β-actin served as the internal control. The sequences of primers we used are listed in **Supplementary Table S3**.

### RNase R and actinomycin D treatment

RNase R and actinomycin D treatments were conducted to validate the stability of circRNA. For RNase R treatment, 1000ng total RNA was incubated with or without 3 U RNase R (Selleck Chemicals, Shanghai, China) at 37 °C for 15 minutes. For actinomycin D treatment, 10 ml/mL actinomycin D (Selleck) was used, and DMSO was used as a negative control. Relative RNA expression was estimated by qRT-PCR.

### Fluorescence in situ hybridization (FISH)

For circRNA FISH, we used a fluorescent in situ hybridization kit (RiboBio, Guangzhou, China). Cells were fixed and permeabilized, then blocked for 30 min at 37 °C with the pre-hybridization buffer. After incubating with a cy3-labeled circNSD1(6) FISH Probe Mix and hybridization buffer overnight at 37 °C, cells were washed with hybridization washing buffer and 0.2 × SSC. The subcellular location of circNSD1(6) was observed with a fluorescent microscope (Olympus, Tokyo, Japan).

### Nuclear-cytoplasmic Fraction

The subcellular location of circNSD1(6) was confirmed by nuclear-cytoplasmic fraction assay. The nucleus and cytoplasm of cells were separated with a PARIS kit (Invitrogen, Carlsbad, USA) under the manufacturer’s instructions.

### RNA intervention and lentiviral infection

For RNA intervention, control siRNA (si-NC), circNSD1(6) siRNA, and linearNSD1 siRNA were synthesized at RiboBio (Guangzhou, China). Lipofectamine 3000 (lipo3000; Invitrogen) and OptiMEM (Gibco) were used to transfect cells.

Stable knockdown of circNSD1(6) was performed by lentiviral infection. Human Lenti-shcircNSD1(6)-GFP (sh-circNSD1(6)) and Lenti-sh-control-GFP (sh-NC) were purchased from Obio Technology (Shanghai, China). Standard procedures were followed to achieve lentiviral infection.

### RNA sequencing

All RNA sequencing was performed by Novogene (Beijing, China). Illumina NEBNext UltraTM RNA Library Prep Kit for Illumina® (NEB, USA) was referred to as sequencing libraries. Transcripts expression with |Log2 FoldChange| >= 1, adjusted P-value<0.05 were filtered, and considered statistically significant.

### Cell proliferation assays

The proliferation ability of cells was accessed by Cell Counting Kit-8 (CCK-8) assays and 5-ethynyl-2’-deoxyuridine (EdU) assays.

CCK-8 assays were performed under the manufacturer’s instructions (Bimake, Houston, TX, USA). Absorbance at 450nm wavelength was measured at 24 h, 48 h, 72 h, 96 h, and 120 h after cells were planted into 96-well plates.

The EdU assay kit was purchased from RiboBio (Guangzhou, China). 8 x 103 cells were seeded in 96-well plates and allowed to adhere for 24 hours. The medium was replaced by the complete medium containing 50μM EdU solution. After incubating for 4 hours, cells were stained with Apollo 567 and 4’,6-diamidino-2-phenylindole (DAPI; Servicebio, Wuhan, China) subsequently. Fluorescent images were visualized with a microscope (Olympus).

### Colony formation assays

The colony formation capability of cells was accessed by colony assays. 1 x 103 cells were seeded in 6-well plates. After incubating for 14 days, cells were stained with 0.5% crystal violet (Servicebio) and counted.

### Apoptosis assays

For apoptosis assays, cells were digested and washed with PBS twice. After resuspending in the binding buffer, cells were stained with annexin V-FITC and PI using the Annexin V-FITC/Propidium Iodide (PI) Apoptosis Detection Kit (BD Biosciences, Franklin Lakes, NJ, USA). Cells were stained for 30 mins and analyzed using a flow cytometer (LSRFortessa X-20; BD).

### Reactive Oxygen Species (ROS) detection

Cellular ROS was detected with a ROS detection probe coupled with green fluorescence (Solarbio, Beijing, China). Cells and H2O2-treated cells were incubated with the ROS detection probe for 20 mins at 37 °C, with the protection of light. The fluorescence was imaged with a fluorescent microscope (Olympus). Besides, the fluorescence can be semi-quantified using flow cytometry. Cells were digested after incubation, and a flow cytometer (LSRFortessa X-20; BD) was utilized to evaluate fluorescent intensity.

### RNA pull-down assays

Biotin-labeled circNSD1(6) (sense) and control (antisense) probes were synthesized by Sangon Biotech (Shanghai, China). RNA pull-down assays were conducted as described^16^. In brief, 2 x 107 cells were washed in ice-cold PBS and subsequently lysed in the washing buffer containing DTT, PMSF, Recombinant Ribonuclease Inhibitor, and protease inhibitor (MCE, NJ, USA). Cell lysates were incubated with 3 μg biotinylated probes for 2 h at 4 °C. A total of 50 μl pre-washed streptavidin magnetic beads (MCE) were added to each binding reaction and further incubated at room temperature for an hour. After washing the beads with a washing buffer six times, Bounded proteins were analyzed by mass spectrometry or western blotting.

### LC-MS/MS analysis

The identification of circRNA binding protein was performed using LC-MS/MS. Proteins were subjected to 10% SDS-PAGE gel. The protein bands that were different from the control were chopped. Peptide mixtures extracted from the chopped gels were identified by LC-MS/MS analysis.

### Western blotting

Protein was extracted with RIPA buffer (Biosharp, Hefei, China) supplemented with PMSF and protease inhibitor (MCE). Western blotting was conducted using standard laboratory western blotting techniques. Polyvinylidene fluoride (PVDF) membranes (Millipore, Darmstadt, Germany) were incubated with primary antibodies at 4 °C overnight, and placed in secondary antibodies diluted with PBST at room temperature for 1 h. Signal detection and image acquisition were realized by Electrochemiluminescence (ECL) luminescent solution (Biosharp) and ChemiDoc XRS + imaging system (BioRad). Primary antibodies used in our study were listed in **Supplementary Table S4**.

### Co-immunoprecipitation

For co-immunoprecipitation (co-IP), cells were lysed with the same procedure using an IP buffer. Cell lysates were pre-washed with protein A/G magnetic beads (MCE), and incubated with primary antibodies on a shaker at 4 °C. The complex was collected, washed with an IP buffer repeatedly, and heated in 2 x SDS Loading Buffer to denaturalize. Western blotting was conducted to verify if target proteins interact with the protein precipitated by primary antibodies.

### The soluble and insoluble protein fraction

Soluble and insoluble proteins were separated using an NP-40-containing lysis buffer. Cells were lysed with an NP-40-containing lysis buffer on ice for 30 mins. After centrifuging at 4 °C for 10 mins, the supernatant (soluble fraction) and pellet (insoluble fraction) were collected separately. Insoluble fractions were redissolved with a lysis buffer containing 1% SDS. Fractions were boiled at 95 °C before undergoing western blotting.

### RNA Immunoprecipitation (RIP)

RNA immunoprecipitation was performed as previously described [3]. Briefly, 2 x 107 cells were washed with ice-cold PBS and lysed in a 200 μL RIP lysis buffer. 50 μL pre-washed protein A/G magnetic beads (MCE) were mixed with 5 μg antibody for 1 h. The protein-magnetic beads mixture was added to the cell lysate and incubated overnight at 4 °C. The next day, beads were washed carefully six times, and RNA was purified with proteinase K buffer. RNA was recovered using Trizol (Vazyme) extraction and analyzed by qPCR.

### Immunofluorescence (IF)

Immunofluorescence assays were performed to observe the location and expression of certain proteins. Cells were seeded in confocal dishes. 24 hours later, cells were fixed and permeabilized, followed by blocking and staining with primary antibodies overnight. A cy3–conjugated goat anti-rabbit IgG (H+L) antibody was used as a secondary antibody, and nuclei were stained with DAPI (Servicebio).

### Autophagosome and autolysosome detection

The autophagosomes and autolysosomes were detected by the RFP-GFP-LC3 reporter system. The stubRFP-sensGFP-LC3 lentivirus was purchased from Genechem (Shanghai,China). Due to the character that GFP fluorescence can be quenched in acidic circumstances, lysosomes are labeled by RFP only. The autophagosomes are reported by the merge of GFP and RFP fluorescence, hence appearing yellow. Therefore, a reporter system of autophagy flux was established. The autophagy flux was observed with a laser scanning confocal microscope (Nikon, Tokyo, Japan).

The autophagosomes and autolysosomes were also directly imaged by transmission electron microscope (TEM). The cells were fixed in 3% glutaraldehyde at 4 °C for 2 h, post-fixed, dehydrated, cut, and stained. Finally, the images were captured using a JEM-1200EX transmission electron microscope (Japan Electronics and Optics Laboratory, Tokyo, Japan).

### Animal experiments

All animal experiments were conducted under the supervision of the Animal Care Committee of Tongji Medical College (IACUC ID: 3360, Hubei, China).

The BALB/c nude mice (4 weeks old, ♀) were purchased from Vital River Laboratory Animal Technology Co. Ltd. (Beijing, China). MCF7-sh-NC and MCF7-sh-circNSD1(6) cells were used to perform human cell line-derived subcutaneous xenograft models. 107 cells were resuspended in serum-free medium and mixed 1:1 with Matrigel (BD, Biosciences). Cells were injected into the axillary flank (10 mice per group). Tumor formation rates for each group were summarized and compared. At the end of the experiment, mice were sacrificed and tumors were excised and measured.

### Immunohistochemistry staining (IHC)

For immunohistochemistry staining, tissues were fixed in 4% paraformaldehyde. Paraffin-embedded sections were prepared and dewaxed. Antigens were unmasked using a citrate buffer. Endogenous peroxidases were depleted with 3% H2O2. Antibodies against p62, Ki-67, and Nrf2 (Proteintech, Wuhan, China) were used. Sections were viewed by a microscope (Olympus), and fields were randomly selected to evaluate the expression of stained markers.

### Statistical analysis

Images were processed and measured with the software ImageJ. The data are displayed as mean ± standard deviation (SD) or as values directly. Grouped two-tailed Student’s t-test was used to assess the significance of the difference. Statistical analyses were conducted using SPSS 24.0 (SPSS Inc. Chicago, IL, USA) and GraphPad Prism 8.0 software (GraphPad Software, San Diego, CA). P value < 0.05 was regarded as statistically significant.

All statistical analyses were performed using R statistical environment (v4.1.0) The type of test method used for statistical analysis was specified in the text where the results were described and details for the test can be found in the relevant figure legend and method section. All tests were two-sided unless otherwise specified.

### Data Analysis & Visualization

Data analysis was conducted using R (v4.1.0) and Python (v3.8.18). Data visualization was conducted using ggplot2 (v3.36)^17^, VennDiagram (v1.7.3)^18^, ComplexUpset (v1.3.3)^19^, ComplexHeatmap (v.2.12.0)^20^, BoutrosLab.plotting.general (v. 7.1.0)^21^, DESeq2 (v1.40.2)^22^, AnnotationDbi (v.1.62.2)^23^, DOSE (v.3.26.2)^24^, org.Hs.eg.db (v.3.17.0)^25^, Seaborn (v.0.12.2)^26^,

Scipy (v1.10.1)^27^, Matplotlib (v3.7.3)^28^. Figure 1A was created using BioRender.com.

### Data Availability

The raw sequencing data of shRNA screening generated in this study can be accessed at NCBI under the accessions GSE269567 (reviewer token: gpadomagbhcdfst). The RNA-seq data of the normal and patient tumors generated in this study have been deposited to GSA database (https://ngdc.cncb.ac.cn/education/tutorials/gsa-human/) under the accession HRA008284.

### Code Availability

All the code used for this analysis is open-source (MIT license).

## Author Contributions

**Designed studies:** W.L, T.L., H.H.H. and M.J.

**Performed experiments:** W.L., F.S., Z.H., X.X., W.Y., M.T.,

**Data Analysis:** T.L., P.H.H., M.C., S.C., Y.Z.

**Manuscript First Draft:** W.L., T.L., P.H.H., M.J and H.H.H.

**Revised & approved manuscript:** All authors

## Acknowledgement

This work was supported by the Science Fund of the National Natural Science Foundation of China (No. 82270830 to J.M.), Hubei Provincial Science & Technology Innovation Team Grant (No. 2022CFB072 to J.M.), Princess Margaret Cancer Foundation (886012001223 to H.H.H.), CIHR operating grants (142246, 152863, 152864 and 159567 to H.H.H.), Terry Fox New Frontiers Program Project Grant (PPG19-1090 and PPG23-1124 to H.H.H.). H.H.H. holds Tier 1 Canada Research Chair in RNA Medicine. M.T. was supported by a CIHR Doctoral Award. P.H.H was supported by a MOHCCN Health Informatics Award and a CIHR Doctoral Award.

## Supplementary Material

### Supplementary Figures

**Supplementary Figure S1.**
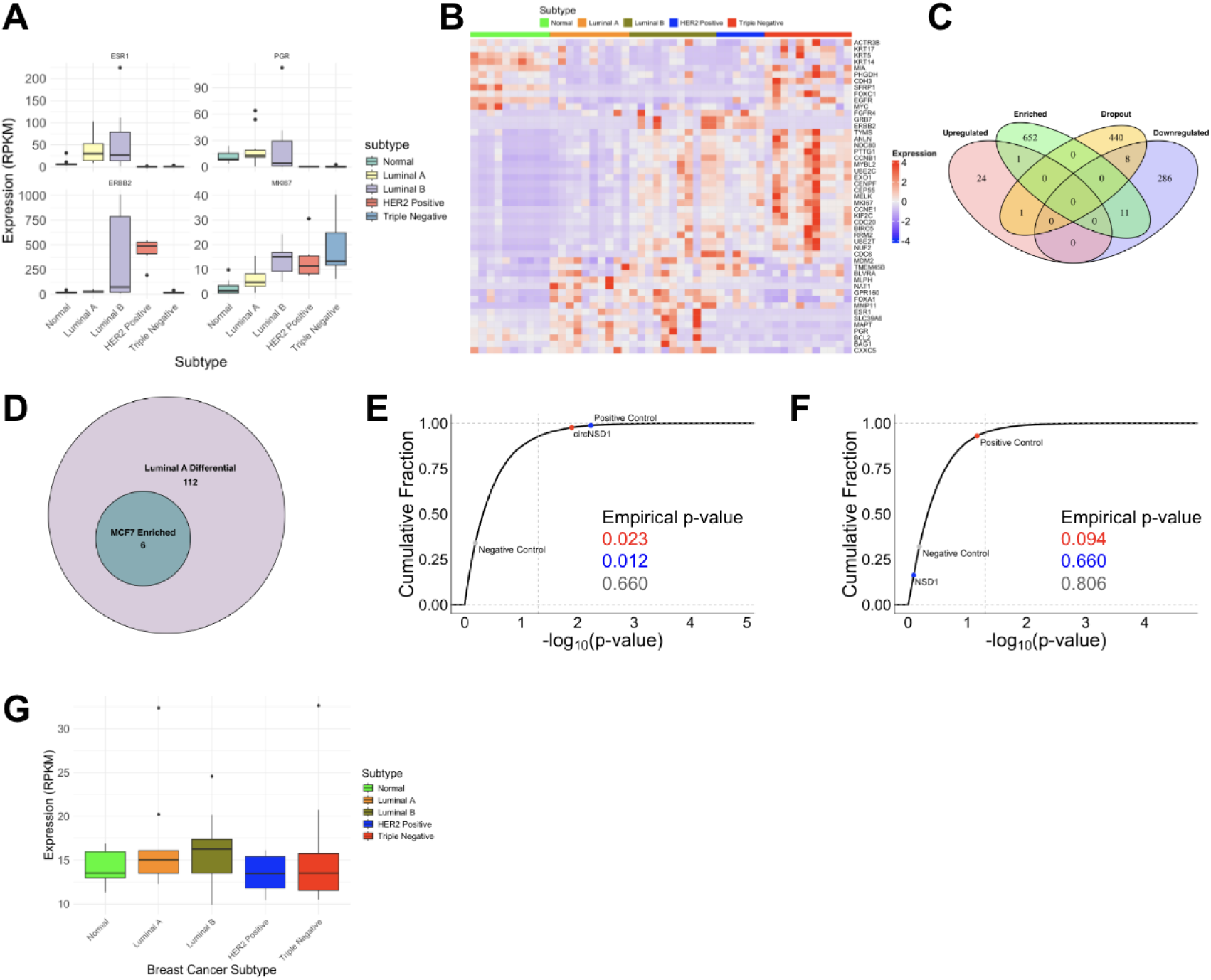
Circular RNA profiling and functional screen in breast cancer. (A) Boxplots showing subtype-specific key genes in breast cancer such as ESR1, PGR, ERBB2 and MKI67. ESR1 Mann Whitney U tests against normal: Luminal A: 3.25×10^-4^, Luminal B: 4.00×10^-4^, HER2 Positive: 2.50×10^-4^, Triple Negative B: 4.00×10^-4^. PGR Mann Whitney U tests against normal: Luminal A: 4.81×10^-1^, Luminal B: 8.09×10^-1^, HER2 Positive: 2.50×10^-4^, Triple Negative B: 5.67×10^-6^. ERBB2 Mann Whitney U tests against normal: Luminal **A**: 7.50×10^-2^, Luminal B: 6.10×10^-2^, HER2 Positive: 2.50×10^-4^, Triple Negative **B**: 7.56×10^-1^. MIKI67 Mann Whitney U tests against normal: Luminal A: 3.50×10^-2^, Luminal B: 6.80×10^-5^, HER2 Positive: 2.00×10^-3^, Triple Negative B: 1.13×10^-5^. (B) PAM50 expression profiles to classify breast cancer subtypes. (C) Venn diagram of upregulated and downregulated circRNAs in Luminal A and dropout and enriched hits from MCF7 circRNA screen. Log2 fold change greater than 1 (for upregulated) and less than −1 (for downregulated) and bonferroni adjusted p-value less than 0.01. (D) Venn diagram of enriched circRNAs from MCF7 screen that overlapped with differential circRNAs in luminal A. Log2 fold change greater than 1 (for upregulated) and less than −1 (for downregulated) and bonferroni adjusted p-value less than 0.01. (E) Cumulative distribution of the negative log_10_(p value) for all circular transcripts. The empirical p-value is derived by applying the ECDF to the log p-value of each point. (F) Cumulative distribution of the negative log_10_(p value) for all linear transcripts. The empirical p-value is derived by applying the ECDF to the log p-value of each point. (G) Expression of NSD1 across breast cancer subtypes.

### Supplementary Tables

**Supplementary Table S1**. Number of Detected circRNAs per Patient.

**Supplementary Table S2**. List of potential circNSD1(6) interacting proteins identified from RNA pulldown mass spectrometry analysis.

**Supplementary Table S3.**
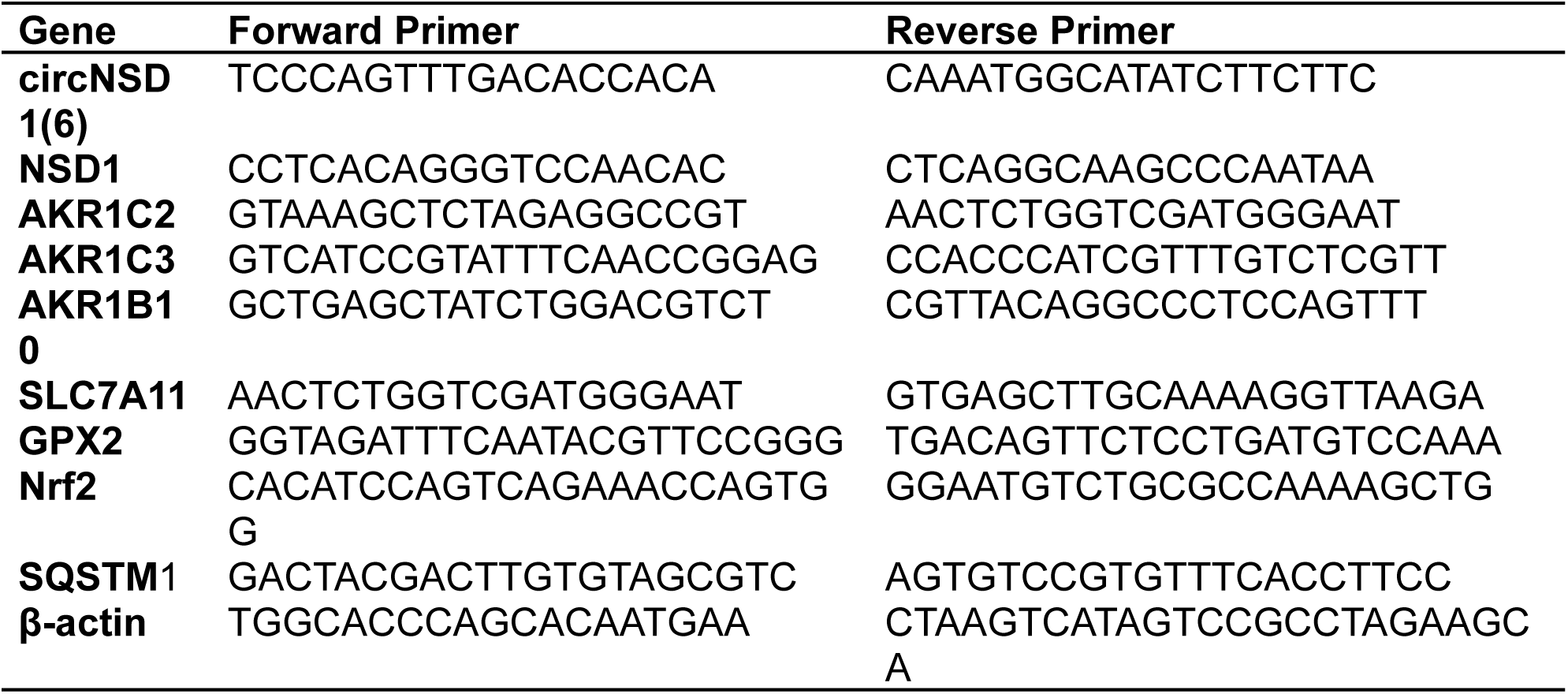
Sequences of primers used in this study.

**Supplementary Table S4.** Primary antibodies used in this study.

| Target | Source | Catalog No. |
| --- | --- | --- |
| p62 | Proteintech | 18420-1-AP |
| Nrf2 | Proteintech | 16396-1-AP |
| LC3 | Proteintech | 14600-1-AP |
| Keap1 | Proteintech | 10503-2-AP |
| Ubiquitin | Cell Signaling Technology | 3936S |
| β-actin | Cell Signaling Technology | 8457S |

## References

1. Sung, H. et al. Global Cancer Statistics 2020: GLOBOCAN Estimates of Incidence and Mortality Worldwide for 36 Cancers in 185 Countries. CA Cancer J. Clin. 71, 209–249 (2021).

2. Loibl, S., Poortmans, P., Morrow, M., Denkert, C. & Curigliano, G. Breast cancer. Lancet 397, (2021).

3. Beilerli, A. et al. Circular RNAs as biomarkers and therapeutic targets in cancer. Semin. Cancer Biol. 83, (2022).

4. Liu, C. X. & Chen, L. L. Circular RNAs: Characterization, cellular roles, and applications. Cell 185, (2022).

5. Kristensen, L. S., Jakobsen, T., Hager, H. & Kjems, J. The emerging roles of circRNAs in cancer and oncology. Nat. Rev. Clin. Oncol. 19, (2022).

6. Moscat, J. & Diaz-Meco, M. T. p62 at the Crossroads of Autophagy, Apoptosis, and Cancer. Cell 137, 1001 (2009).

7. Moscat, J., Karin, M. & Diaz-Meco, M. T. p62 in Cancer: Signaling Adaptor Beyond Autophagy. Cell 167, (2016).

8. Harris, I. S. & DeNicola, G. M. The Complex Interplay between Antioxidants and ROS in Cancer. Trends Cell Biol. 30, (2020).

9. de la Vega M, R., Chapman, E. & Zhang, D. D. NRF2 and the Hallmarks of Cancer. Cancer Cell 34, (2018).

10. Jena, K. K. et al. TRIM16 controls assembly and degradation of protein aggregates by modulating the p62-NRF2 axis and autophagy. EMBO J. 37, (2018).

11. Komatsu, M. et al. The selective autophagy substrate p62 activates the stress responsive transcription factor Nrf2 through inactivation of Keap1. Nat. Cell Biol. 12, (2010).

12. Ma, X.-K., Xue, W., Chen, L.-L. & Yang, L. CIRCexplorer pipelines for circRNA annotation and quantification from non-polyadenylated RNA-seq datasets. Methods 196, 3–10 (2021).

13. Feng, X. et al. Myosin 1D and the branched actin network control the condensation of p62 bodies. Cell Res. 32, 659–669 (2022).

14. Bjørkøy, G., Lamark, T. & Johansen, T. p62/SQSTM1: a missing link between protein aggregates and the autophagy machinery. Autophagy 2, 138–139 (2006).

15. Jain, A. et al. p62/SQSTM1 is a target gene for transcription factor NRF2 and creates a positive feedback loop by inducing antioxidant response element-driven gene transcription. J. Biol. Chem. 285, 22576–22591 (2010).

16. Nielsen, A. F. et al. Best practice standards for circular RNA research. Nat. Methods 19, (2022).

17. Wickham, H. ggplot2: Elegant Graphics for Data Analysis. (Springer, 2016).

18. Chen, H. & Boutros, P. C. VennDiagram: a package for the generation of highly-customizable Venn and Euler diagrams in R. BMC Bioinformatics 12, 35 (2011).

19. Lex, A., Gehlenborg, N., Strobelt, H., Vuillemot, R. & Pfister, H. UpSet: Visualization of Intersecting Sets. IEEE Trans. Vis. Comput. Graph. 20, 1983–1992 (2014).

20. Gu, Z., Eils, R. & Schlesner, M. Complex heatmaps reveal patterns and correlations in multidimensional genomic data. Bioinformatics 32, 2847–2849 (2016).

21. P’ng, C. et al. BPG: Seamless, automated and interactive visualization of scientific data. BMC Bioinformatics 20, 42 (2019).

22. Love, M. I., Huber, W. & Anders, S. Moderated estimation of fold change and dispersion for RNA-seq data with DESeq2. Genome Biol. 15, 550 (2014).

23. AnnotationDbi. Bioconductor https://bioconductor.org/packages/AnnotationDbi.

24. Yu, G., Wang, L.-G., Yan, G.-R. & He, Q.-Y. DOSE: an R/Bioconductor package for disease ontology semantic and enrichment analysis. Bioinformatics 31, 608–609 (2015).

25. Carlson, M. org.Hs.eg.db. (Bioconductor, 2017). doi:10.18129/B9.BIOC.ORG.HS.EG.DB.

26. Waskom, M. seaborn: statistical data visualization. J. Open Source Softw. 6, 3021 (2021).

27. Virtanen, P. et al. SciPy 1.0: fundamental algorithms for scientific computing in Python. Nat. Methods 17, 261–272 (2020).

28. Hunter, J. D. Matplotlib: A 2D Graphics Environment. Comput. Sci. Eng. 9, 90–95 (May-June 2007).

